# Re-Annotator: Annotation Pipeline for Microarrays

**DOI:** 10.1101/019596

**Authors:** Janine Arloth, Daniel Magnus Bader, Simone Röh, Andre Altmann

**Affiliations:** Translational Research Department, Max Planck Institute of Psychiatry, Kraepelinstrasse 2-10, 80804, Munich, Germany; Gene Center Munich, Ludwig-Maximillians-Universität München, Feodor-Lynen Strasse 25, 81377, Munich, Germany; Department of Neurology and Neurological Sciences, Stanford University, School of Medicine, 780 Welch Road, CJ350 C38, CA-94304 Palo Alto, USA

## Abstract

**Background:** Microarray technologies are established approaches for high throughput gene expression, methylation and genotyping analysis. An accurate mapping of the array probes is essential to generate reliable biological findings. Manufacturers typically provide incomplete and outdated annotation tables, which often rely on older genome and transcriptome versions differing substantially from up-to-date sequence databases.

**Results:** Here, we present the Re-Annotator, a re-annotation pipeline for microarrays. It is primarily designed for gene expression microarrays but can be adapted to other types of microarrays. The Re-Annotator is based on a custom-built mRNA reference, used to identify the positions of gene expression array probe sequences. A comparison of our re-annotation of the Human-HT12-v4 microarray to the manufacturer’s annotation led to over 25% differently interpreted probes.

**Conclusions:** A thorough re-annotation of probe information is crucial to any microarray analysis. The Re-Annotator pipeline consists of Perl and Shell scripts, freely available at http://sourceforge.net/projects/reannotator. Re-annotation files for Illumina microarrays Human HT-12 v3/v4 and MouseRef-8 v2 are available as well.

## Background

Analysis of gene expression profiles under various conditions is one of the corner stones in modern molecular biology research. One major challenge in working with gene expression microarrays is the quality of the annotation of the array probes used by the platform. Differences in probe annotations complicate the replication of studies as well as metaanalyses across platforms. Moreover, the annotations provided by the manufacturers quickly become outdated with every update of the genome assemblies as well as the accompanying annotation tables. For example, the number of annotated transcripts in the RefSeq Gene database (RefSeq release 59 [1]) differs from hg18 (NCBI build 36.1) to hg19 (GRCh37 build 37) assembly by more than 1,200 transcripts (43,236 to 44,596). Furthermore, the initial mapping provided by the manufacturers contains several severe problems – some probes map to non-transcribed genomic regions, bind secondary targets or have other properties that may confound a proper analysis, such as common SNPs in the probe sequence. Clearly, these probes should be removed from the analysis as an accurate probe annotation is fundamental for all downstream analyses and ensures accurate biological interpretation of the results. Outdated annotation of probes becomes an increasing problem in publicly available gene expression catalogs such as the ALLEN brain atlas [2] as researchers tend to use the provided expression data as is, that is, without further validity checks. In order to identify probes with potential annotation problems, a sound re-annotation of all probes is required. Recently, a small number of approaches were developed that allow the reannotation of gene expression microarray data by re-aligning the probe sequences to the entire human genome [3, 4]. By using the whole genome as the mapping reference, the likelihood of short reads to be mapped to multiple locations and intergenic regions increases, thereby decreasing the number of uniquely mappable probes [5]. Still 24% of the human genome cannot be uniquely mapped using 50 bp long sequences with two mismatches [6, 7], which corresponds to the sequence length of Illumina array probes. Therefore, including untranscribed regions, which theoretically cannot even be part of the cRNA library of interest, reduces the mappability and introduces an additional source of unnecessary errors. Our approach considers these mappability issues; we developed the Re-Annotator pipeline for gene expression microarrays that directly maps probes to a custom-built mRNA reference. Therefore our pipeline enables us to correctly annotate various formerly non-mappable probes [5].

## Implementation

### Requirements

Prior installation requirements include the Burrows-Wheeler Aligner (BWA) [8], SAMtools [9], ANNOVAR [10] and Perl. Data requirements include the genome assembly of interest, a corresponding gene annotation table, microarray probe sequences and optionally SNP data (e.g., dbSNP).

### Creating the *in silico* mRNA reference

The sequences of each transcript (exons only) are extracted and concatenated to serve as an mRNA reference (see Figure 1).

**Figure 1:**
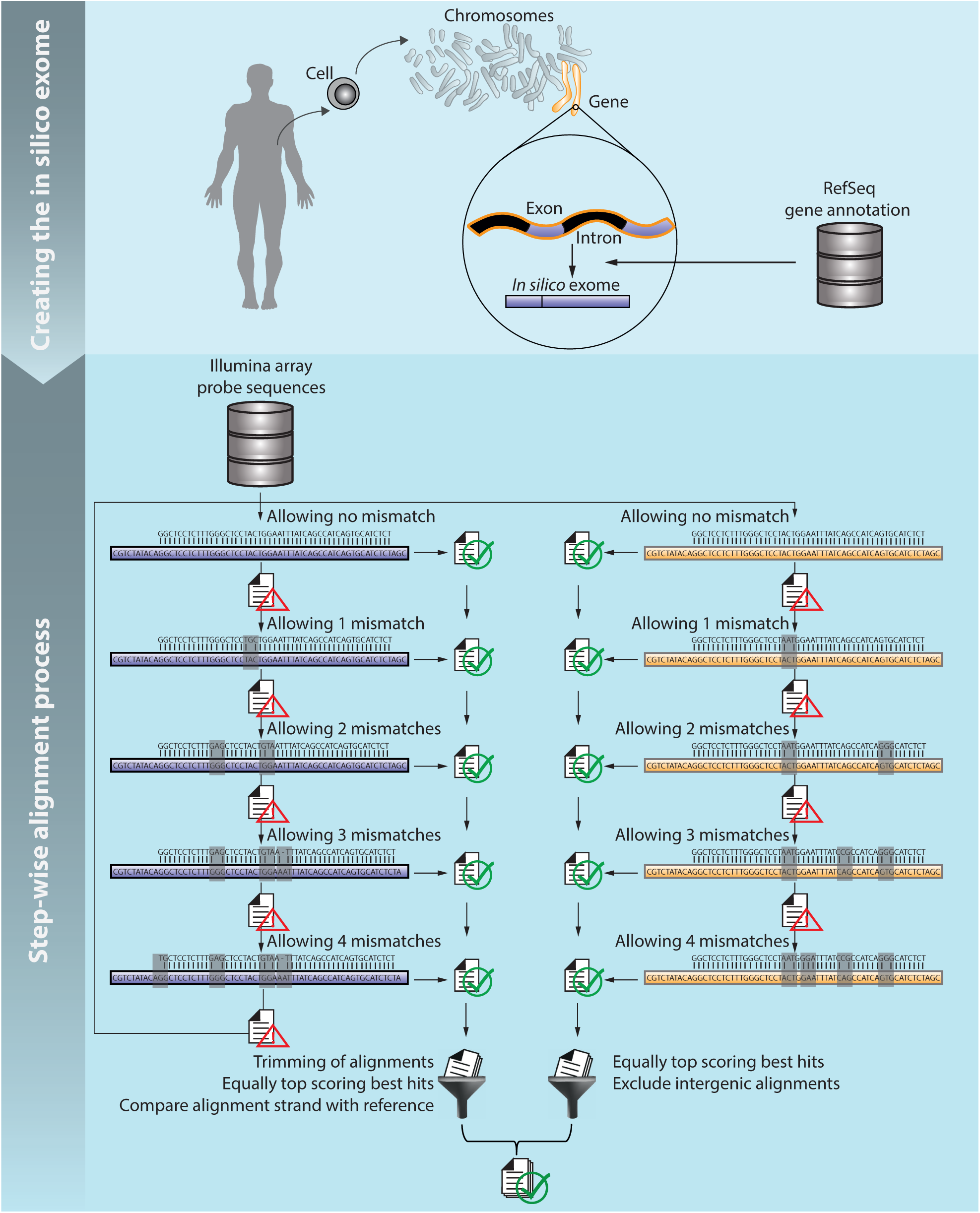
Annotation pipeline. Schematics of the computational pipeline flow: (1) creating the *in silico* reference and (2) step-wise alignment process. Purple sequences in the left column correspond to alignments to the *in silico* reference and brown sequences in the right column correspond to alignments to the genome.

### Stepwise alignment process

The annotation process starts with the alignment of the probe sequences provided by the manufacturer to the mRNA reference using BWA. The quality of the probes does not decrease by position, thus the aligner is executed without seeding (seed length *>* length of probe sequence). A step-wise alignment process is performed allowing no mismatches at first. All unaligned probe sequences are resubmitted for alignment with the number of allowed mismatches increased by one. Array probes that do not align to the mRNA reference with a limit of four mismatches are mapped to the whole genome using the same step-wise approach. Alignments are stored in SAM format and all alternative best hits are included. The alignment results are processed according to the following steps:

### mRNA reference alignments

1. alignments starting or ending with a mismatch, insertion or deletion are trimmed
2. alignment positions are mapped to the genome
3. alignments are discarded if the alignment strand does not match the original coding strand information

### Genomic alignments

1. step (1) and (3) of mRNA reference alignments procedure
2. gene annotation is added using ANNOVAR
3. annotations mapping to intergenic or up-and downstream regions (defined as 1 kb distance to the transcription start or end site) are excluded

### SNP information

Optionally, SNP information can be added to the annotation.

### Post processing

In order to ensure that a probe is specific for one genomic region, we eliminate probes with multiple hits that cannot be assigned to the same gene, i.e., we recommend that coordinates of different hits should be not more than 25 bp apart (default for Illumina arrays) from each other. This threshold is largely arbitrary (half of the probe length of Illumina probes) and it may remove relevant probes, but it will guarantee that the probes are aligned uniquely to a distinct region. We require that probes with multiple annotations should be aligned in the same direction (alternate haplotype regions of the original assembly were ignored). The final re-annotation is provided with additional information on the probe gene symbol and updated position (also for splice-annotations).

## Results

### Re-Annotator improves annotation of human probes compared to manufacturer

We analyzed the Illumina HumanHT-12 v4 probe sequences (*n* = 47,230) using the Re-Annotator Pipeline. 95% of the array probes (see Figure 2a and Table 1) were aligned to either our mRNA reference (*n* = 34,277) or to the genome (*n* = 10,661). Probes aligned to genomic locations without any known transcribed gene (*n* = 7,493) were excluded from the further re-annotation process. After the post-processing filter, 77.7% of all aligned probes (see Table 1) mapped to a distinct region (defined as a maximum of 25 bp distance between the hits) in the genome and were included in the final annotation file for the HumanHT-12 v4 BeadChip array (referred to as reliable array probes in the following). The majority (93.8%) of those reliable probes (see Figure 2b and Table 1) were aligned without mismatches. The number of hits per probe to a region ranged from 1 to 32, where 67.7% had only one unique hit and 96% (*n* = 33,539) had less than five hits (see Figure 2c). The vast majority of uniquely mapped probes (92.1%, 32,160 out of 34,936; see Figure 2d and Table 1) resided in regions without known SNPs in the Caucasian population (provided by the 1,000 Genomes Project; see methods section). It is conceivable that SNPs within the probe sequence may be the source of “differential” expression via altered hybridization efficiency. However, Schurmann et al. [4] reported no consistent effects of SNPs located in array probes on hybridization efficiency. Thus, one has to test individually whether these SNPs are associated with alternate expression signals intensity. Roughly 23.5% (*n* = 11,086) of all Illumina probes were not annotated with probe coordinates and gene symbols by the manufacturer. About 36,6% (*n* = 4,062) of these unannotated probes could be rescued and reliably re-annotated. For 21.5% (*n* = 7,789) of the probe sequences with a complete Illumina annotation (*n* = 36,144) the Re-Annotator provides an annotation that differs from the manufacturer’s annotation.

**Figure 2:**
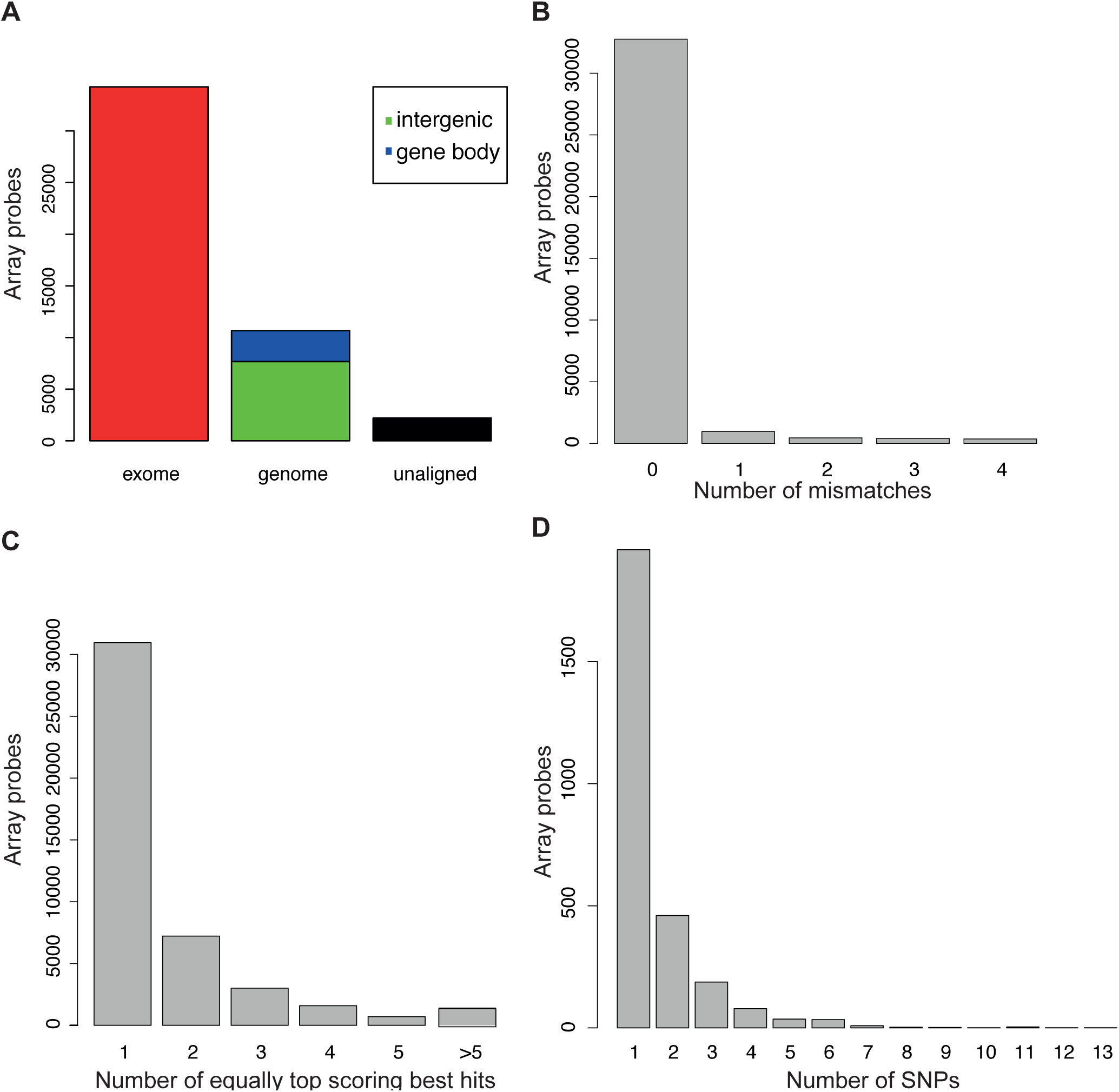
Results of the re-annotation of Illumina probe sequences for the HumanHT-12 v4 expression BeadChips. A) Barplot of the alignment basis. The red bar represents the array probe sequences that could be aligned to the *in silico* build reference. The bar in the middle represents those sequences, which could only be aligned to the genome while the black bar represents sequences, which could not be aligned at all with our criteria. Sequences aligning to the genome where further differenced whether they map to genes or intergenic regions. Histograms of distribution of B) mismatches within array probe sequences, C) the number of equally top scoring best hits per array probe sequence and D) the number of SNPs within each array probes sequences.

**Table 1:**
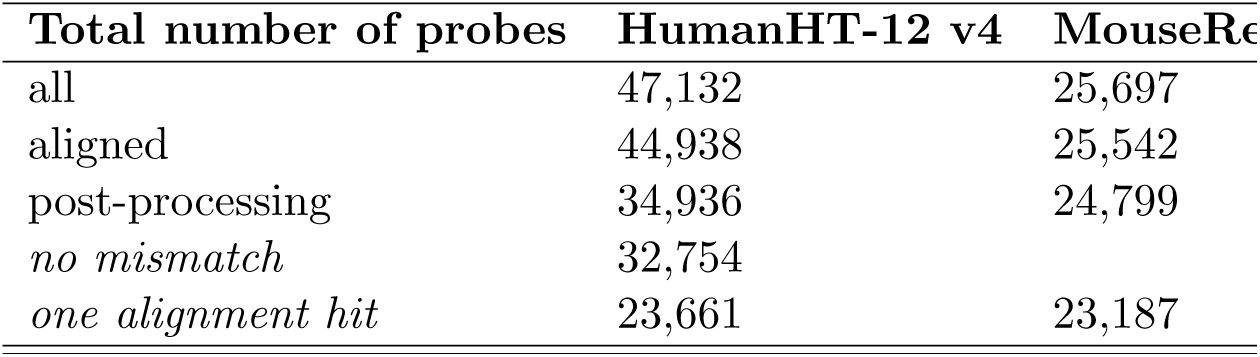
Results of re-annotation of Illumina probe sequences.

### Re-Annotator provides probes with less variable intensity then ReMOAT

Comparing Re-Annotator to ReMOAT [3], another re-annotation tool, we found that 86.3% (*n* = 29,759) of probes annotated as “reliable” by ReMOAT (quality equal to “Perfect” and “Good”; *n* = 34,476 transcripts) were also classified as reliable probes by Re-Annotator (see Figure 3a). However, 5% (*n* = 1,532) of these 29,795 array probes received different annotations by the two tools. A total of 5,177 probes, which were annotated as reliable by ReMOAT, were excluded by Re-Annotator due to alignments in intergenic regions (63.7%), multiple different hits in the mRNA reference (26%) and no alignment (10.3%).

**Figure 3:**
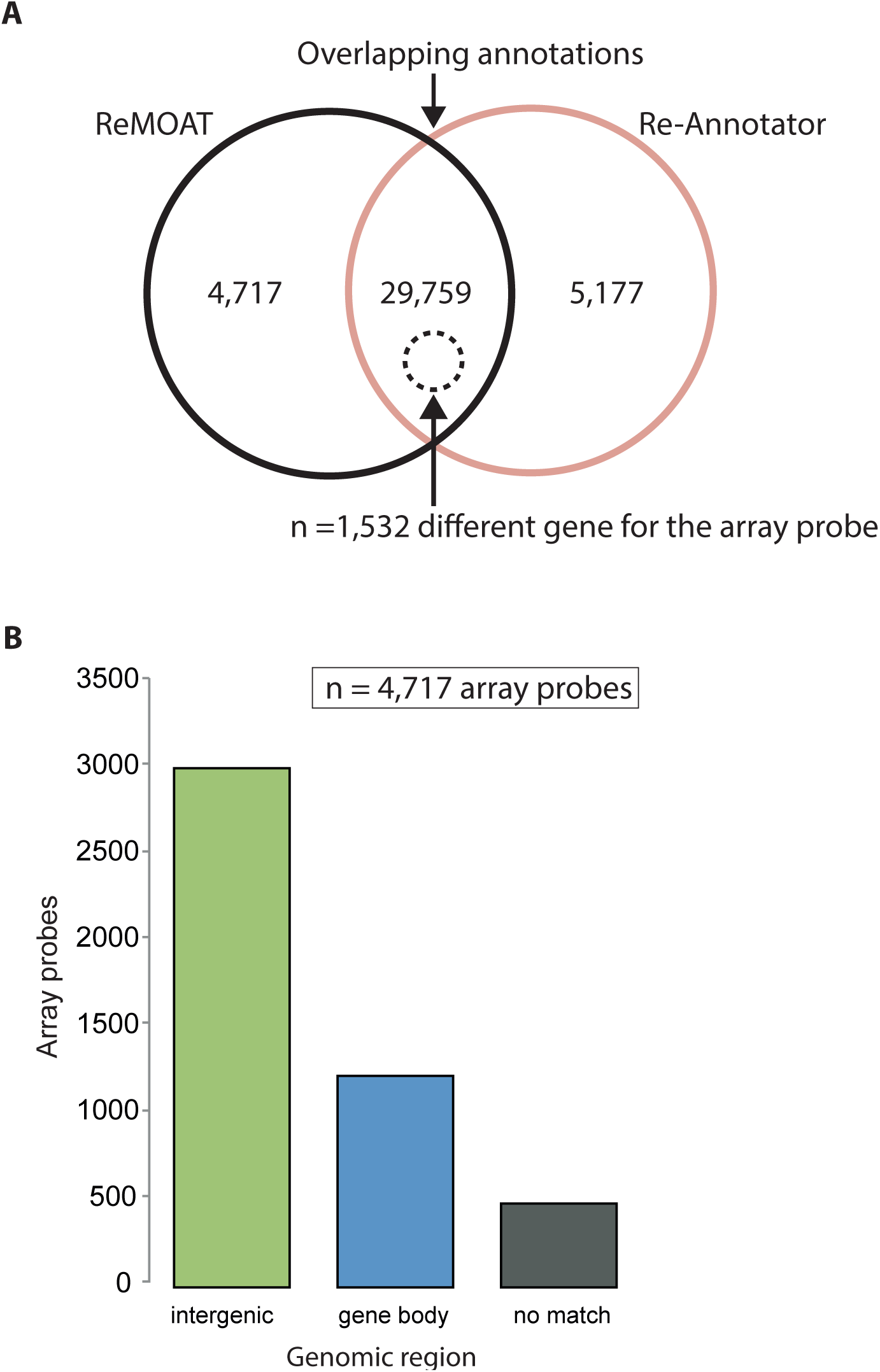
Comparison of our re-annotation of Illumina probe sequences for the HumanHT-12 v4 expression BeadChips and a previous re-annotation. A) Venn diagram representing the overlap of transcripts annotated with ReMOAT (black circle) and the Re-Annotator (red circle). 5% (black dashed circle) of the overlapping annotations map to different genes. B) The bar graph showing the distribution of microarray probe sequences excluded from our re-annotation.

In order to further demonstrate the advantage of Re-Annotator, we reanalyzed gene expression profiles in whole blood cells from 36 individuals at baseline (without stimulation) hybridized to the Illumina Human HT-12 v4.0 microarray chips (Gene Expression Omnibus GSE64930, see Additional File 1). Assuming that probes with a bad annotation capture more noise from the data and thus will exhibit a higher average variance than the remaining probes with good annotation quality, we sought to quantify this effect in a real-world dataset. Briefly, we created five groupings:

1. probes in repetitive regions identified by ReMOAT using RepeatMasker (*n*=1,176)
2. probes excluded by Re-Annotator but included by ReMOAT (*n*=433)
3. probes included by Re-Annotator but excluded by ReMOAT (*n*=1,208)
4. probes excluded by both Re-Annotator and by ReMOAT (*n*=377)
5. probes included by both Re-Annotator and by ReMOAT (*n*=9,236)

Next, we computed the variance for each probe across all 36 subjects and compared the mean variance of the probes in groups 1-3 to the mean variance of the probes in groups 4 and 5. In group 5 we randomly selected probes to match the numbers in groups 1-3. Compared to group 4 (0.124 *±* 0.14), group 3 (0.109 *±* 0.12; Wilcoxon P=0.011) but not group 2 (0.132 *±* 0.37; P=0.094) showed a significantly reduced mean probe variance. Further, as expected probes with good annotations in both tools (group 5) showed the lowest variance (*≈* 0.09), with group 2 showing a significant increase in variance (P=7.2 *×* 10^-6^) that was not observed in group 3 (P=0.25). ReMOAT also excludes probes that are located in repeats as detected by RepeatMasker. Our experiment showed that probes in repeat regions appear rather normal with significantly reduced variance (0.099 *±* 0.093) compared to group 4 (P=0.016) and unchanged variance compared to group 5 (0.095 *±* 0.1; P=0.13).

In addition to analyzing the variance of probes across samples, using the real-world data, we compared the expression status of the probe to the annotation (include vs exclude) provided by both tools. Here we hypothesized that probes that were excluded were less likely to pass the detection threshold for expression. As expected, probes that were annotated as “bad” were less likely to be expressed, the effect was significantly stronger (P*<*0.05) for Re-Annotator (OR: 3.9 95% CI: 3.65-4.16) than for ReMOAT (OR: 2.11 95% CI: 2.01-2.23).

### Re-Annotator refines annotation of mouse probes compared to manufacturer

Additionally, we analyzed the Illumina MouseRef-8 v2 probe sequences (*n* = 25,697) using our Re-Annotator pipeline. Almost all probes were aligned (99.4%; see Figure 4a and Table 1) to either the mRNA reference (*n* = 24,994) or genome (*n* = 548) and passed the post-processing filter (see Table 1). 97.9% of all post-processed array probes were aligned without mismatches (see Figure 4b and Table 1) and 93.5% mapped to a single region with only 14 probes having 5 or more hits per probe (see Figure 4c and Table 1). The successful re-annotation of the mouse microarray can be explained by the reduced array content to only Ref-Seq genes, providing good transcriptomic annotation quality.

**Figure 4:**
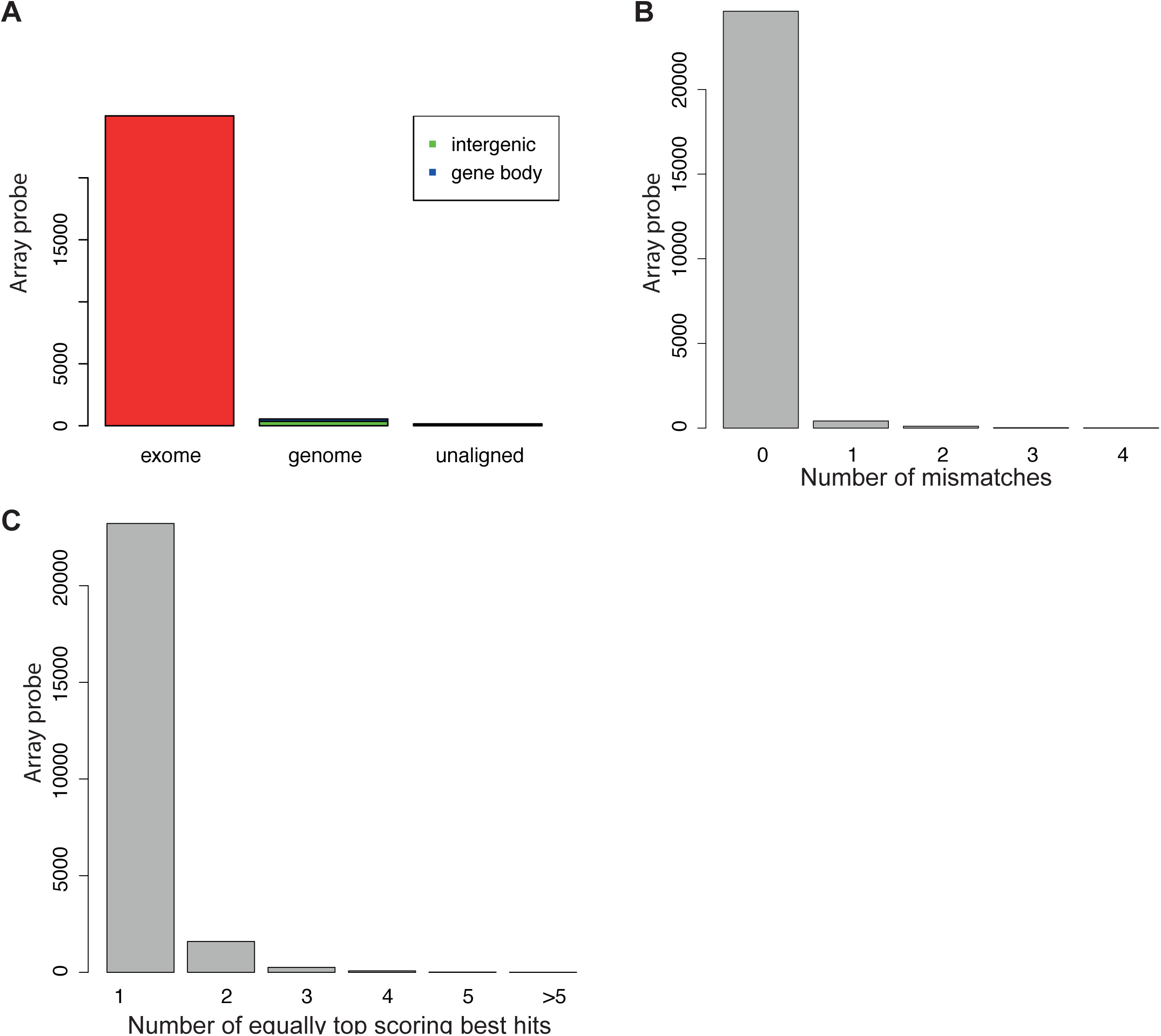
Results of the re-annotation of Illumina probe sequences for the MouseRef-8 v2 expression BeadChips. A) Barplot of the alignment basis. The red bar represents the array probe sequences that could be aligned to the *in silico* build reference. The middle bar represents those sequences, which could only be aligned to the genome while the black bar represents sequences, which could not be aligned at all. Sequences aligning to the genome where further differenced whether they map to genes or intergenic regions. Histograms of distribution of B) mismatches within array probe sequences and C) the number of equally top scoring best hits per array probe sequence.

We further provide an annotation file for Illumina Human HT-12 v3 and a file including the overlapping content of both human Illumina arrays (Illumina HT-12 v3 and v4).

## Discussion

A precise annotation of microarray probe sequences is essential for accurate biological findings and replicability. In this work, we present a pipeline to reannotate probe sequences of gene expression microarrays using a custom-built mRNA reference and applied it to three Illumina BeadChip arrays (Human HT-12 v3, v4 and MouseRef-8 v2). The re-annotation revealed that indeed numerous array probes were incompletely (23.5%) or incorrectly (16.5%) annotated by the manufacturer. A source of such mis-annotation may be due to changes in genome assembly or changes in exon/intron boundaries since the original design of the chip.

Over 21% of re-annotated probes were assigned to different genes as given by the manufacturer. For example, three of the five Illumina HT-12 v4 array probe sequences illustrated in Figure 5, all perfectly re-annotated within the first or second exon of the human gene *ISCA1* on chromosome 9 using the Re-Annotator (see Figure 5), were originally annotated on chromosome 5 (see Figure 5) within an intergenic region. A reason for such a discrepancy could be that the probe sequences were designed using an older assembly version (hg18). When using the old version the probes target a gene at the specified location, which was rearranged in the more recent assembly (hg19). Hence, it is important to keep the annotation tables of the probes up-to-date. ReMOAT, a previous re-annotation [3], based on a genomic alignment, placed these probes in accordance with our annotation (see Figure 5). We recommend to check all given probe sequence annotations (second matches as well as other given genomic matches), also when using the ReMOAT annotation, as the given genomic location might be wrong. Such an example is illustrated in Figure 6; the probe sequence was allocated to an intergenic region (see Figure 6). We annotated this probe sequence to be on chromosome 17 within an exon of *ABCA9*, which was in accordance with the second match of the ReMOAT annotation.

**Figure 5:**
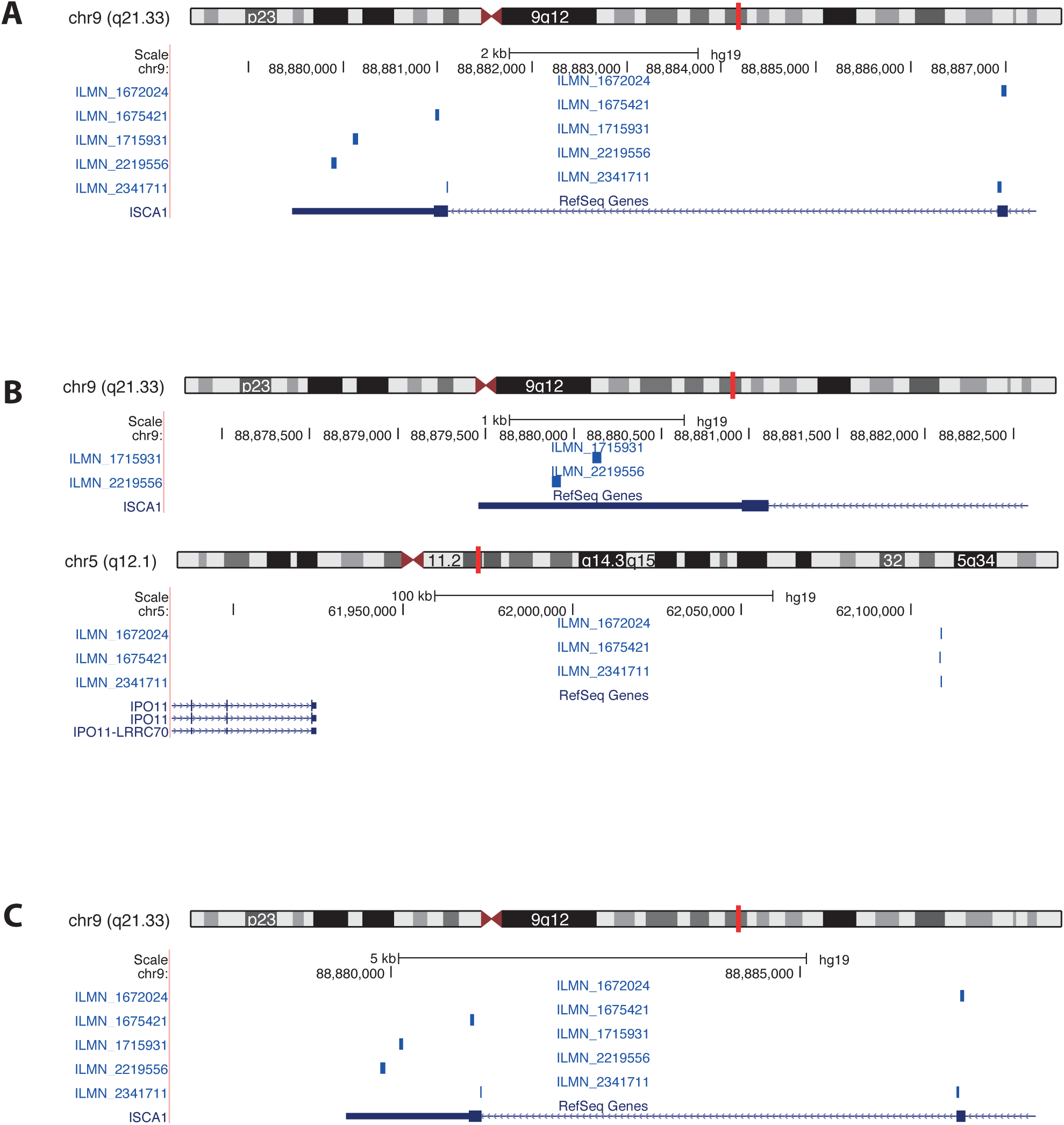
UCSC genome browser graphic for the human *ISCA1* gene on chromosome 9, including the targeting Illumina probes (ILMN 1715931, ILMN 1672024, ILMN 2219556, ILMN 1675421 and ILMN 2341711). Custom tracks representing the probe sequences annotated by A) the Re-Annotator, B) manufacturer and C) ReMOAT.

**Figure 6:**
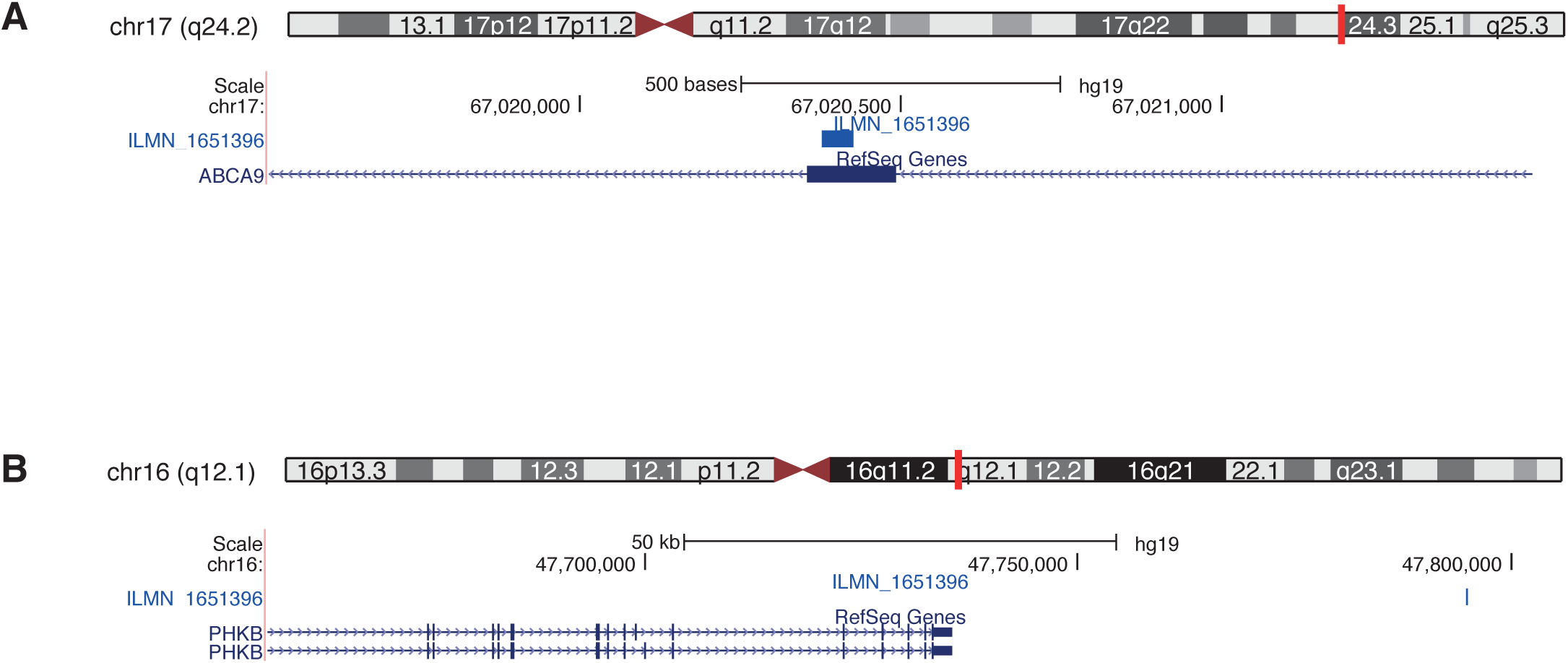
UCSC genome browser graphic for the human *ABCA9* gene on chromosome 9, including the targeting Illumina probe (ILMN 1651396). Custom tracks representing the probe sequences annotated by A) the Re-Annotator and B) manufacturer and ReMOAT.

Furthermore, the Re-Annotator does no filtering based on the RepeatMasker^1^ as suggested by Barbosa-Morais et al. [3]. Many regions marked by this algorithm are in fact perfectly mappable, and filtering may eliminate data on important genes. An example is a probe located within the *FKBP5* gene (ILNM 1778444). If repeats are masked, this probe is marked as unreliable since it is located within a short interspersed nuclear element (SINE). Still, there is no issue of mappability or uniqueness, the probe should not be excluded from further analysis. To further proof the reliability of probes removed on the basis of RepeatMasker, we compared the variance of those probes to probes annotated as unreliable (bad quality and excluded in both Re-Annotator and ReMOAT) and found that Repeat-Masker excluded probes show a lower variance than bad probes.

Approximately 74% of all human probes present on the latest Illumina gene expression array (HT-12 v4) were uniquely allocated to one gene locus. Such a re-annotation is important for removing uninformative probes, such as probes that cannot be placed into a distinct region, before starting differential gene expression analysis. This increases specificity of an analysis and will decrease the false discovery rate. With our pipeline we closed these gaps and compensated for wrong annotations.

## Conclusion

A thorough re-annotation of probe sequences is not a standard part of gene expression microarray analysis. To highlight its profound effect, we applied our pipeline to Illumina BeadChip Human HT-12 v4 and compared it to the Illumina annotation as well as to the ReMOAT annotation. We discovered that the Human HT-12 v4 re-annotation differs significantly (by 25% and 16%, respectively) from the annotations provided by Illumina and ReMOAT Our pipeline improves the probe annotation and proves to be an essential step in producing high quality microarray results.

## Availability and requirements

### Software

Project name: Re-Annotator

Project homepage: http://sourceforge.net/projects/reannotator

Operating system: Linux and Mac OSX

Program language: Perl and Shell scripts

Other requirements: BWA, SAMtools and ANNOVAR (see instructions in zip package)

License: open-source

### Application data

For the re-annotation of human microarrays, we used UCSC version 19 of the human genome^2^. We obtained the RefSeq transcript coordinates^3^ (download on January 2012). Similarly, for the reannotation of mouse microarrays, we downloaded the July 2007 assembly of the mouse genome (mm9) from the UCSC Genome Browser^4^ and the Ref-Seq transcript coordinates^5^ (download on January 2012). We obtained Illumina annotations^6^ and downloaded the results of a previous re-annotation [11]. The gene expression profiles in whole blood cells from 36 individuals at baseline (without stimulation) hybridized to the Illumina Human HT-12 v4.0 microarray chips were downloaded from Gene Expression Omnibus (GSE64930).

## Competing interests

The authors declare that they have no competing interests.

## Author’s contributions

AA and DMB developed this software. JA and SR tested the software intensively. JA is responsible for data analysis. JA and SR wrote the initial version of the manuscript. AA and DMB edited the manuscript. The manuscript has been seen and approved by all authors.

## Acknowledgments

We would like to thank Peter Weber for useful discussions. DMB is supported by a DFG Fellowship through the Graduate School of Quantitative Biosciences Munich (QBM).

## Additional Files

Additional File 1: Additional Methods and Analysis

## Footnotes

1 www.repeatmasker.org

2 http://hgdownload.soe.ucsc.edu/goldenPath/hg19/bigZips/chromFa.tar.gz

3 http://hgdownload.soe.ucsc.edu/goldenPath/hg19/database/refGene.txt.gz

4 http://hgdownload.cse.ucsc.edu/goldenPath/mm9/bigZips/chromFa.tar.gz

5 http://hgdownload.cse.ucsc.edu/goldenPath/mm9/database/refGene.txt.gz

6 http://www.switchtoi.com/annotationfiles.ilmn

## References

[1] Pruitt, K.D., Tatusova, T., Brown, G.R., Maglott, D.R.: NCBI Reference Sequences (RefSeq): current status, new features and genome annotation policy. Nucleic Acids Research 40(Database issue), 130–5 (2012)

[2] Hawrylycz, M.J., Lein, E.S., Guillozet-Bongaarts, A.L., Shen, E.H., Ng, L., Miller, J.A., van de Lagemaat, L.N., Smith, K.A., Ebbert, A., Riley, Z.L., Abajian, C., Beck-mann C.F., Bernard, A., Bertagnolli, D., Boe, A.F., Cartagena, P.M., Chakravarty, M.M., Chapin, M., Chong, J., Dalley, R.A., Daly, B.D., Dang, C., Datta, S., Dee, N., Dol-beare T.A., Faber, V., Feng, D., Fowler, D.R., Goldy, J., Gregor, B.W., Haradon, Z., Haynor, D.R., Hohmann, J.G., Horvath, S., Howard, R.E., Jeromin, A., Jochim, J.M., Kinnunen, M., Lau, C., Lazarz, E.T., Lee, C., Lemon, T.A., Li, L., Li, Y., Morris, J.A., Overly, C.C., Parker, P.D., Parry, S.E., Reding, M., Royall, J.J., Schulkin, J., Sequeira, P.A., Slaughterbeck, C.R., Smith, S.C., Sodt, A.J., Sunkin, S.M., Swanson, B.E., Vawter, M.P., Williams, D., Wohnoutka, P., Zielke, H.R., Geschwind, D.H., Hof, P.R., Smith, S.M., Koch, C., Grant, S.G.N., Jones, A.R.: An anatomically comprehensive atlas of the adult human brain transcriptome. Nature 489(7416), 391–399 (2013)

[3] Barbosa-Morais, N.L., Dunning, M.J., Samara-jiwa A.S., Darot, J.F.J., Ritchie, M.E., Lynch, A.G., Tavaré, S.: A re-annotation pipeline for Illumina BeadArrays: improving the interpretation of gene expression data. Nucleic Acids Research 38 (2010)

[4] Schurmann, C., Heim, K., Schillert, A., Blankenberg, S., Carstensen, M., Dörr, M., Endlich, K., Felix, S.B., Gieger, C., Grallert, H., Herder, C., Hoffmann, W., Homuth, G., Illig, T., Kruppa, J., Meitinger, T., Müller, C., Nauck, M., Peters, A., Rettig, R., Roden, M., Strauch, K., Völker, U., Völzke, H., Wahl, S., Wallaschofski, H., Wild, P.S., Zeller, T., Teumer, A., Prokisch, H., Ziegler, A.: Analyzing illumina gene expression microarray data from different tissues: methodological aspects of data analysis in the metaxpress consortium. PLoS one 7(12), 50938 (2012)

[5] Lee, H., Schatz, M.C.: Genomic dark matter: the reliability of short read mapping illustrated by the genome mappability score. Bioinformatics 28(16), 2097–2105 (2012)

[6] Koehler, R., Issac, H., Cloonan, N., Grim-mond S.M.: The uniqueome: a mappability resource for short-tag sequencing. Bioinformatics 27(2), 272–274 (2011)

[7] Derrien, T., Estellè, J., Marco Sola S., Knowles, D.G., Raineri, E., Guigo, R., Ribeca, P.: Fast computation and applications of genome mappability. PLoS one 7(1), 30377 (2012)

[8] Li, H., Durbin, R.: Fast and accurate short read alignment with Burrows-Wheeler transform. Bioinformatics 25(14), 1754–1760 (2009)

[9] Li, H., Handsaker, B., Wysoker, A., Fennell, T., Ruan, J., Homer, N., Marth, G., Abeca-sis G., Durbin, R., 1000 Genome Project Data Processing Subgroup: The Sequence Alignment/Map format and SAMtools. Bioinformatics 25(16), 2078–2079 (2009)

[10] Wang, K., Li, M., Hakonarson, H.: ANNOVAR: functional annotation of genetic variants from high-throughput sequencing data. Nucleic Acids Research 38(16), 164 (2010)

[11] Dunning, M., Lynch, A., Eldridge, M.: illuminaHumanv4.db: Illumina HumanHT12v4 Annotation Data (chip illuminaHumanv4)

